# Efficient gradient boosting for prognostic biomarker discovery

**DOI:** 10.1101/2021.07.06.451263

**Authors:** Kaiqiao Li, Sijie Yao, Zhenyu Zhang, Biwei Cao, Christopher M. Wilson, Pei Fen Kuan, Ruoqing Zhu, Xuefeng Wang

**Author notes:** These authors contributed equally to this work (Kaiqiao Li, Sijie Yao, Zhenyu Zhang).

## Abstract

**Motivation:** Gradient boosting decision tree (GBDT) is a powerful ensemble machine learning method that has the potential to accelerate biomarker discovery from high-dimensional molecular data. Recent algorithmic advances, such as Extreme Gradient Boosting (XGB) and Light Gradient Boosting (LGB), have rendered the GBDT training more efficient, scalable and accurate. These modern techniques, however, have not yet been widely adopted in biomarkers discovery based on patient survival data, which are key clinical outcomes or endpoints in cancer studies.

**Results:** In this paper, we present a new R package **Xsurv** as an integrated solution which applies two modern GBDT training framework namely, XGB and LGB, for the modeling of censored survival outcomes. Based on a comprehensive set of simulations, we benchmark the new approaches against traditional methods including the stepwise Cox regression model and the original gradient boosting function implemented in the package **gbm**. We also demonstrate the application of **Xsurv** in analyzing a melanoma methylation dataset. Together, these results suggest that **Xsurv** is a useful and computationally viable tool for screening a large number of prognostic candidate biomarkers, which may facilitate cancer translational and clinical research.

**Availability:** **Xsurv** is freely available as an R package at: https://github.com/topycyao/Xsurv

**Supplementary information:** Supplementary data are available at *Bioinformatics* online.

## 1 Introduction

Boosting is one of the most powerful and successful techniques introduced in the fields of statistics and machine learning for solving complex classification and regression problems. The basic idea is to sequentially learn a combination of multiple weak learners to create a more sophisticated learning model that achieves better prediction performance. AdaBoost, developed in 1995 (Freund and Schapire, 1997), is the first practical implementation in this category. In early 2000’s, a more flexible and effective solution called gradient boosting machine (GBM) was proposed by Friedman (Friedman, 2001). GBM generalizes the boosting idea to any loss function which is differentiable. By combining the gradient descent algorithm, GBM in each stage identifies an add-on weak learner function by fitting on the negative gradient of the loss function (the “pseudo-residuals”). The main reasons for the widespread application of GBM today are because of its flexibility, extensibility and easy implementation. It works for various popular loss functions and can be coupled with different types of weak learners from simple regression to decision trees (i.e. GBDT). Extensions of GBM have been implemented and integrated in various R packages, most notably the gbm (Greenwell, et al., 2007) and **caret** package (Kuhn, 2020). Nonetheless, GBM faces several drawbacks that might lead to inferior prediction performance in real data analysis. First, like other greedy-search-based algorithms, GBM and GBDT can converge to local optimal and tends to overfit the training data, especially with a large learning rate and more iterations. Second, GBM involves more hyperparameters than methods such as random forest (uses two tuning parameters), making hyper-parameter tuning more challenging and producing less reproducible results in practice.

Extreme gradient boosting (Chen and Guestrin, 2016), or XGBoost (XGB), is a recent advancement that builds on the GBDT framework (Mason, et al., 1999) (Hastie, et al., 2009) and has rapidly gained considerable prominence in the field of applied machine learning. Because of its superior and reliable predictive performance across a variety of test data sets, XGB has become a de facto benchmark algorithm in data science competitions (e.g. Kaggle) and real-world machine learning projects. XGB significantly mitigates the overfitting issue by introducing extra regularization, built-in tree pruning, and subsampling features. XGB is especially attractive for its computational efficiency. As will be further discussed in the method section below, the optimization problem in tree boosting is greatly simplified by using a trick in calculating the gain score that can be extended to a customized loss function. In addition, it leverages multithreading parallel computing offered by modern CPUs. Therefore, this framework scales well to both large sample size and feature numbers that conventional packages like gbm are not geared to handle.

LightGBM (Jeschke, et al., 2017), hereafter referred to as LGB, is another significantly improved gradient boosting tree algorithm that has achieved great popularity recently. It integrates multiple novel techniques to jointly optimize computation speed, memory usage and prediction performance. The most notable differences compared to other gradient boosting methods are that LGB uses the leaf-wise growth strategy (instead of level-wise strategy) to construct the tree and it adopts the gradient-based one-side sampling (GOSS) to find a split. GOSS allows a more efficient (and thus “lighter”) implementation of GBDT, which keeps all the data with large gradients and performs random sampling on the data with small gradients. In addition, LGB combined GOSS with a new algorithm called exclusive feature bundling (EFB) to reduce number of features and further improve efficiency. When the training data set contains ultra-high dimensional features or extremely large sample size, LGB is regarded as a more suitable alternative to XGB and other existing gradient boosting methods. The main scope of this study is to investigate the feasibility of modern gradient boosting methods, i.e., XGB and LGB frameworks, for the modeling and prediction of censored survival outcomes. Our work was primarily motivated by a growing demand for more efficient and effective machine learning methods for prognostic biomarker identification in cancer research. Although XGB and LGB provide the interface for customizing objective (loss) function, it is less straightforward to incorporate survival data. A special form of loss function and prediction evaluation metric need to be defined in the context of XGB and LGB workflow because most survival outcomes contain at least two variables, measuring the survival time and censoring information, respectively. A simple approximation is to force the data to a binary classification problem by ignoring the censoring and dichotomizing the time variable, so off-the-shelf machine learning packages can be directly applied. However, it is known that excluding incomplete patient information will result in an inefficient and biased estimation of coefficients in the survival model. This study will focus on building the survival objective function using the Cox proportion hazard (Cox PH) model (Cox, 1972) because it is the widely used in biomedical and health fields. A key characteristic of the objective function based on Cox PH is that it is differentiable and guaranteed to be convex. Of note, the newest version of R package xgboost (Chen, et al., 2021) also allows for the analysis of survival data but only implemented one type of loss function and evaluation metric (based on the Cox likelihood), and specific technical details have not been clearly documented. This paper will review and discuss the implementation of both XGB and LGB for survival data, with the concordance index (or C-index) as a more robust alternative to the Cox loss function.

The remainder of the paper is organized as follows. In Section 2, we briefly describe the efficient boosting method and provides solutions for Cox’s model, as well as the boosting step for boosting the concordance index. In Section 3, we present simulation results to evaluate the predictive performance of the XGB and LGB survival routines and compare the approaches with the standard Cox regression model and gradient boosting method implemented in **gbm**. Finally, we apply the developed survival boosting to a melanoma methylation dataset with the goal to identify targeted CpG sites with prognostic values, followed by discussions.

## 2 Methods

### 2.1 Gradient boosting overview

Similar to other supervised machine learning approach, the ultimate goal of boosting is to find an optimal function of covariates *f*\*(*x*) to predict the outcome *Y*, by minimizing the loss function ℒ(*Y, f*(*x*)). Boosting builds the final predictive model by iteratively combining weak learners that predict the outcome based on a simple model. Thus, it provides an alternative path to build generalized additive models (GAMs): *f*(*x*) = ∑_*j*_ *f*_*j*_(*x*), in which each *f*_*j*_(*x*) is a (weighted) weak learner. Boosting performs variable selection implicitly and works properly even with strong multicollinearity and high dimensionality. By incorporating weighting scheme into resampling steps, boosting is able to focus on more training-informative samples in each step. In the boosted linear regression case, the reweighting is achieved by refitting residuals calculated from previous steps as a surrogate outcome. Gradient boosting generalizes AdaBoost to any smooth loss function ℒ(.,.). In each iteration, gradient boosting trains the base learners by refitting the negative gradients of the loss function. The gradient in this context can be viewed as pseudo-residual. In the special case of square loss, the gradients equal the residuals.

#### 2.1.1 Cox’s survival model for generalized gradient boosting

In biomedical research, the Cox proportional hazard model (Cox 1972) is the most commonly used model for the regression analysis of survival outcomes. In the following, we consider the standard time-to-event (e.g. to death) data for the *i*-th instance (*t*_*i*_, ***x***_*i*_, *δ*_*i*_), where *t*_*i*_ is the observed survival time, ***x***_***i***_ is the covariate vector, and *δ*_*i*_ is the censoring indicator. Censoring occurs when a time-to-event is not observed during follow-up. We focus on the right censoring, in which the actual event time is no earlier than the observed time. The Cox model defines the hazard function of a subject at time *t* to be the product of a baseline hazard *λ*_0_(*t*) and an exponential function of the covariates,

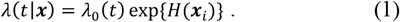

Here *H*(⋅) is a risk score function that relates covariates and regression coefficients. In the standard Cox model, the risk score is a linear term, i.e.,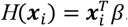. In the case of high-dimensional and non-parametric setting, it is often difficult to specify the functional form of *H*(⋅) explicitly. In boosting framework, the target function *H* is a linear combination of base functions. For most of the following discussion, we will take derivative with respect to the risk term *H*, instead of the original parameters *β*. A key advantage of the Cox model and its partial likelihood (PL) is that the estimation of *β* does not depend on *λ*_0_(*t*), *PL* = ∏{exp (*H*(*x*))/ ∑_*k*∈*R*_ exp (*H*(*x*))}^*δ*^. Here *R* is the set of the observations at risk at time *t*. The key to understanding this formula is to recognize that the *PL* is similar to the conditional probability that a particular study subject is the one that has an event at time *t*. Here we want to minimize the negative of the Cox’s log-partial likelihood as the loss function,

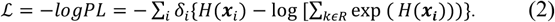

The regression coefficients can be thus estimated by 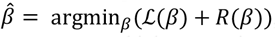, where *R*(*β*) denotes the regularization terms which constrain coefficients, such as the L1/L2 or group lasso penalty terms (Yuan and Lin, 2006) (Simon, et al., 2013). Since the Cox objective function is convex, the problem can be efficiently solved by gradient or subgradient based algorithms.

Both XGB and LGB start with computing the negative gradient direction of the loss function (working/pseudo response or residual) with respect to the current estimate of risk score. The gradient for the Cox’s loss function is thus,

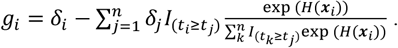

The standard gradient boosting will iteratively optimize the loss function (2) by choosing a weak learner (based on a single or few predictors) that is closest to the negative gradient directions, e.g., at step *m*, the optimal basis function *η*^(*m*)^ can be calculated by

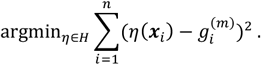

Modern gradient boosting frameworks further consider the second order Taylor expansion of the loss, which basically can be viewed as improved variants of Newton boosting. To simplify the expression, we denote 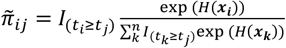. *π* can be treated as estimated absolute probability of failure for subject *i* at time *t*_*j*_ (given that a failure occurs at *t*_*j*_). The empirical gradient function can now be written as

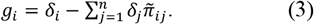

As detailed in Supplementary Materials, it can be shown that the second derivative of the Cox PL loss (with respect to *H*) is

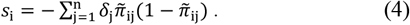

Note that both *g*_*i*_ and *s*_*i*_ are empirical terms evaluated at the given data points. At *m*-th step, we can thus approximate the Cox loss function with functions (3) and (4) through second order Taylor’s series expansion of the loss around the current function *H*^(*m*−1)^,

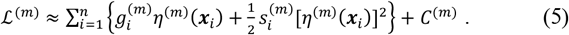

*C* only involves loss term evaluated at *H*^(*m*−1)^, which thus can be treated as a constant term at the current iteration. The above equation provides a new optimization objective, which is equivalent to weighted least square regression problem. By rearranging terms in (5), the optimal basis function can be expressed as

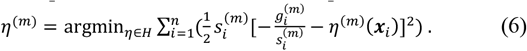

The updated function after step *m* is then *H*^(*m*)^ = *H*^(*m*−1)^ + *vη*^(*m*)^, where parameter *v* represents the learning rate or step size. The step size moving along the selected direction is determined in line search step through fitting a linear proportional hazard model(Li and Luan, 2005).The reason that the new formulation is much more efficient than directly optimizing the original loss function in (2) is that most terms are the same for a given iteration and only need to calculate once, while only the term *η*(***x***_*i*_) need to be evaluated for each candidate function or new split. The computation is particularly efficient when using regression tree as base learner, which will be discussed in the following section.

#### 2.1.2 Efficient tree boosting framework

Tree boosting as proposed originally by (Friedman, 2001) uses decision tress as base learners. Each leaf (terminal) node in a decision tree is assigned with a prediction value or leaf weight. The tree basis function is defined as 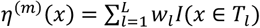, where *w*_*l*_ is the leaf weight and the leaf node indicator function *I*(*x* ∈ *T*_*l*_) defines the structure of a proposed tree. By plugging the tree basis function into (5), the empirical loss function can be rewritten as

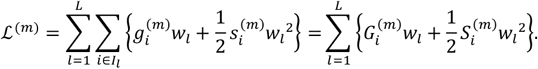

where *G*_*i*_ and *S*_*i*_ are the sum of *g*_*i*_ and *s*_*i*_ in one leaf node. Solving the quadratic function of *w*_*l*_, the optimal solution is thus 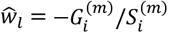. The optimal score function is thus 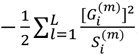. In XGB (Chen and Guestrin, 2016) or LGB (Jeschke, et al.), regularization terms are further incorporated into the loss function to control for model complexity. For example, if the *l*2 regularization term of the leaf weights is considered, the optimal score function becomes 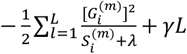, where *λ* is the *l*2 regularization term of the leaf weights and *γ* is the penalty term for the number of terminal nodes. The loss reduction after one split, also known as *gain* score, is given by

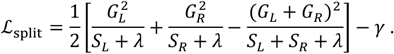

The *l*2 regularization term will not only shrink the leaf weight but also, together with the variance term (*S*), will alter the structure of the final tree. Therefore, the optimization problem is greatly simplified by searching splits that minimize the empirical loss— based on the derived gain function. In the descriptions below, we will use XGB-Cox and LGB-Cox to stand for XGB and LGB based algorithms for solving Cox partial likelihood. The current framework can be easily implemented in R and Python and also allows other customized loss function for survival data as long as it is twice differentiable, such as the smoothed concordance index to be introduced.

#### 2.1.3 Hyperparameter tuning

Similar to XGB and LGB, there are three groups of hyperparameters needed to be determined in **Xsurv**. The first group is basic parameters defining boosting types (e.g., choice of decision tree or linear model) and parameters for basic computational environment. The second group is booster parameters such as the number of trees, tree depth, learning rate, regularization terms, etc. Xsurv provides functions to automatically perform cross-validation to identify these hyperparameters. The default learning rate and regularization terms are set from 0.01 to 0.5, with two parameter search strategies (grid search and random search) offered in the main function options. The third group include more specific learning task parameters that defines the objective function and evaluation metrics for validation data.

### 2.2 Directly boosting the concordance Index

One limitation of the Cox model is the assumption of proportion hazards (2). In Xsurv, we also implement an alternative approach based on boosting the C-index (Harrell, et al., 1982) directly. The C-index is defined as

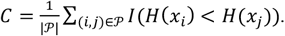

where 𝒫 is the set of orderable pairs and *t*_*i*_ < *t*_*j*_, *H*(⋅) is the assigned risk score. Since the C-index is not differentiable, we use a smoothed concordance index as proposed previously (Chen, et al., 2013),

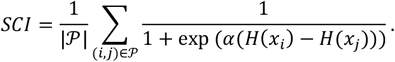

in which *α* is a hyperparameter that controls the steepness. To implement XGB and LGB, we need the first and second derivative of SCI with respect to *H*(⋅). The first derivative of C-index loss function is

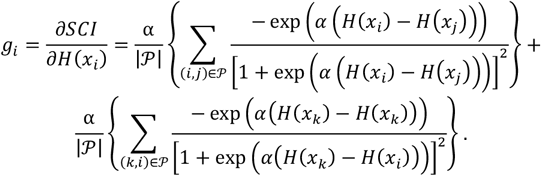

The derivation of second derivative is provided in **Supplementary Materials**. With these two derivatives, we can optimize the loss function in same way as in the Cox model. In the following sections, we denote the C-index based boosting methods as XGB-C and LGB-C.

### 2.3 Survival outcome calibration

One shortcoming of Cox and C-index based boosting methods is that the predicted risk scores are only meaningful at the population level. The predicted risk score cannot be immediately interpreted as survival time or probability of individual patients. To facilitate its usage in the personalized medicine setting, Xsurv provides a function to transform predicted scores back to survival time. In addition, we provide an option to output single-patient prognostic group classification (high, medium and low-risk groups), which are more interpretable as illustrated by the example below.

## 3 Results

In this section, we first present simulation studies that evaluate the performance of the survival gradient boosting methods in terms of risk function estimation and variable selection. We consider survival outcome data generated based on both linear and nonlinear different risk score functions, under three different scenarios. We then illustrate the application of Xsurv with an example of discovering prognostic biomarkers in melanoma using methylation data.

### 3.1 Simulation Scenario 1 (linear model)

In this scenario, we start with a simple proportional hazard model with linear link function. Let covariates *X* = *X*_1_, …, *X*_*p*_ be i.i.d. standard normal distributed random variables and the dimension *p* = 100. The failure time *T* follows an exponential distribution with mean at exp(*X*_1_ + *X*_2_ + … *X*_10_). The censoring time *C* follows an exponential distribution with mean equals 10, in order to prevent tail region values from dominating the prediction error, observations exceeding 4 are forced to be censored.

To benchmark the prediction performance of different algorithms, the following six different methods were conducted: Step-wise Cox(Draper and Smith, 1981; Efroymson, 1960; Hocking, 1976), GBM-Cox, XGB-Cox, XGB-C, LGB-Cox and LGB-C. Two metrics are selected to verify our results: C-index and the Integrated Brier Score (IBS) (Brier, 1950). A total of 100 independent replicates were generated from the same model above, and the total sample size *n* is 1000 in each replicate. The first 800 samples were used as the training data and the remaining 200 as test data. The resulted C-index and IBS are reported in Table 1. As expected, in this simulation setting, Stepwise Cox achieved the best results because it fits the true model. XGB and LGB outperformed GBM with relatively higher C-index.

**Table 1.** Comparisons of predictive performance of the Cox regression, GBM, XGB and LGB based on three simulation scenarios.

| Model | C-index (S.E.) | IBS (S.E.) |
| --- | --- | --- |
| Simulation scenario 1 |  |  |
| Stepwise-Cox | .883 (.009) | .022 (.007) |
| GBM-Cox | .728 (.026) | .054 (.014) |
| XGB-Cox | .782 (.017) | .052 (.010) |
| XGB-C | .798 (.017) | .045 (.010) |
| LGB-Cox | .770 (.018) | .052 (.010) |
| LGB-C | .779 (.018) | .049 (.011) |
| Simulation scenario 2 |  |  |
| Stepwise-Cox | .587 (.023) | .038 (.014) |
| GBM-Cox | .647 (.026) | .036 (.013) |
| XGB-Cox | .704 (.021) | .033 (.012) |
| XGB-C | .704 (.023) | .032 (.012) |
| LGB-Cox | .698 (.026) | .032 (.012) |
| LGB-C | .702 (.024) | .032 (.012) |
| Simulation scenario 3 |  |  |
| Stepwise-Cox | .588 (.021) | .039 (.015) |
| GBM-Cox | .638 (.027) | .037 (.014) |
| XGB-Cox | .702 (.020) | .034 (.013) |
| XGB-C | .702 (.021) | .033 (.013) |
| LGB-Cox | .718 (.020) | .032 (.012) |
| LGB-C | .708 (.021) | .033 (.013) |

### 3.2 Simulation scenarios 2 and 3 (nonlinear models)

In the second scenario we consider nonlinear effects. Let *X* = *X*_1_, …, *X*_*p*_ be i.i.d. standard normal distributed random variables. We simulated 100 replicates, each with a feature dimension *p* =100 and the sample size of 1000. The failure time *T* follows an exponential distribution with mean equals to

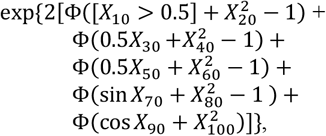

where Φ is the standard normal cumulative distribution function. The censoring time *C* has probability 1/3 to be 0.02 and probability 2/3 to be uniform (0, 0.02), the censoring rate in this case is approximately 30%.

As summarized in Table 1, Stepwise-Cox yielded the worst results. XGB and LGB models still showed satisfactory results given the complicated nonlinear generative model. In order to evaluate the algorithms in terms of feature selection accuracy, we examined the top ranked biomarkers based on the feature importance score from all replicates (Figure 1a). It can be shown that nonlinear signals like *X*_20_, *X*_40_, *X*_60_, *X*_80_ had more chances to be detected by XGB and LGB based survival models. Meanwhile, GBM and the stepwise model selected more null or false positive signals if based on the top-ranking feature list.

**Fig.1.**
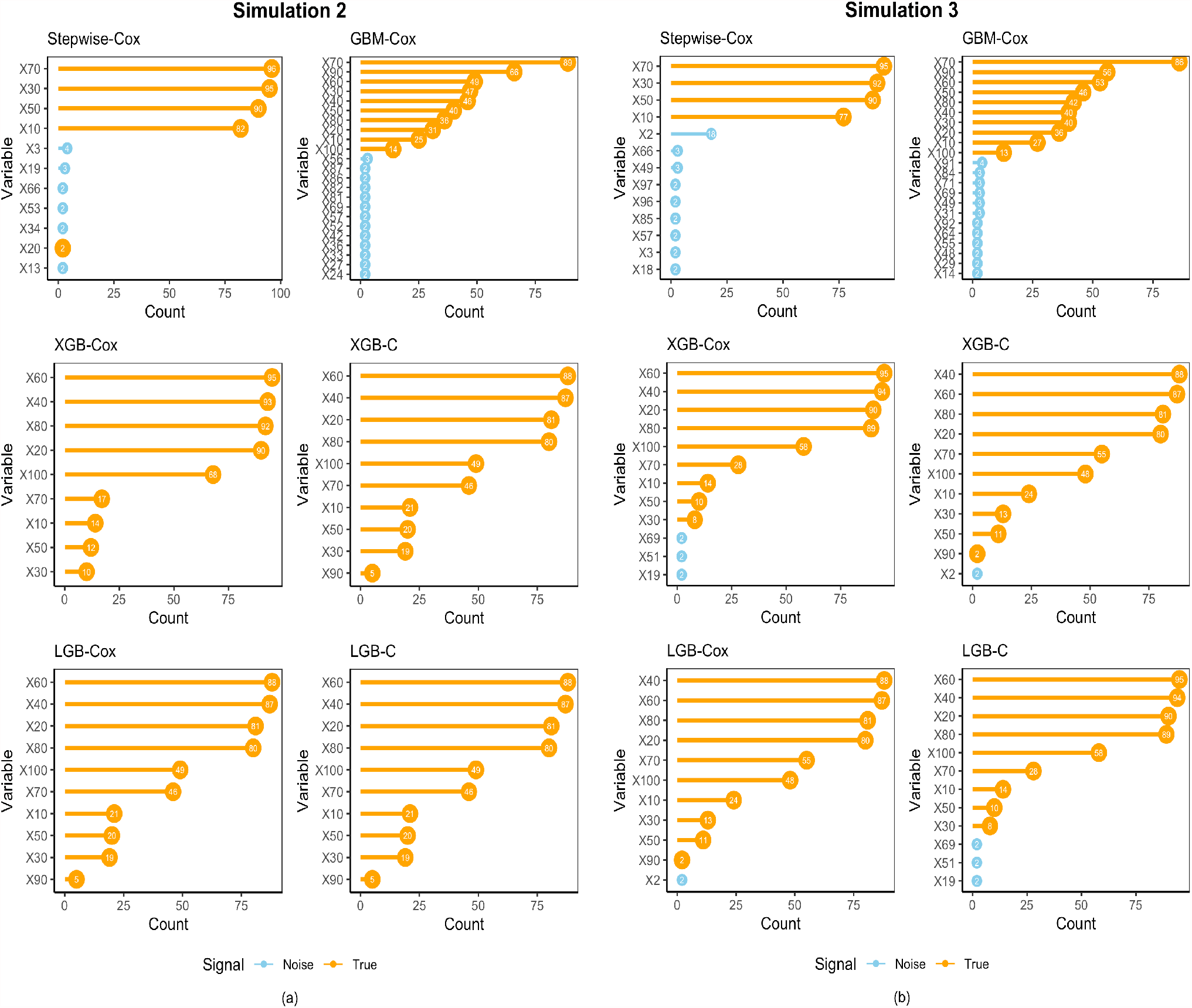
Comparisons of top predictive features selected based on 100 replicates in Simulation 2 (a) and Simulation 3 (b). The y-axes list all variables that appeared in top five features in the final predictive model of each training while x-axes represent the frequencies of being selected. The selected “True” signal features are represented by orange bars, and the selected “Noise” features are represented by blue ones.

In the third simulation scenario, we further considered correlated covariates. This simulation was an extension of the scenario 2 but with covariates X = (X_1_, …, X_100_) following a multivariate normal distribution with covariance matrix V, where V_ij_ = ρ^|i−j|^ and we set *ρ* = 0.5 in this scenario. The results are summarized in Table 1 and Figure 1b. Similar to the previous scenario, overall the new gradient boosting models implemented in Xsurv outperformed the baseline models in terms of overall prediction accuracy and feature selection.

### 3.3 Survival outcome calibration

We further generated a simulation setting to demonstrate the survival calibration function offered by the **Xsurv** package. In this simulation, 100 covariates were generated from a normal distribution with variance 1 and three groups of mean *μ*_*i*_: *μ*_1_ = 0, *μ*_2_ = 10 and *μ*_3_ = 15. We generated 400 samples in each group, hence the total sample size is 1200. The failure time T follows a Weibull distribution with the shape parameter equaling 2 and the scale parameter

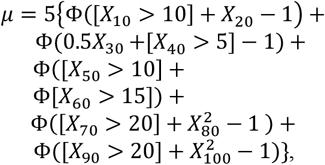

where Φ is the standard normal cumulative distribution function. The censoring time *C* has probability 1/3 to be 0.005 and probability 2/3 to be uniform (0, 0.005). The censoring rate in this case is approximately 30%. For each replicate, we use 1000 samples as training data and the remain 200 as test data. The risk level is defined based on the value of *ν*. A larger value of *μ* is equivalent to a higher risk of mortality. The real risk level for 1200 samples is defined by its corresponding true *ν*. For each replicate, samples were divided into three groups based on tertiles of *ν*, representing “Low Risk”, “Medium Risk”, and “High Risk”, respectively. The result in Figure 2 (based on five replicates) shows that XGB models can successfully classify patients into three risk groups (accuracy around 95%). Similar results were found for LGB models (**Supplementary Fig. 1)**.

**Fig.2.**
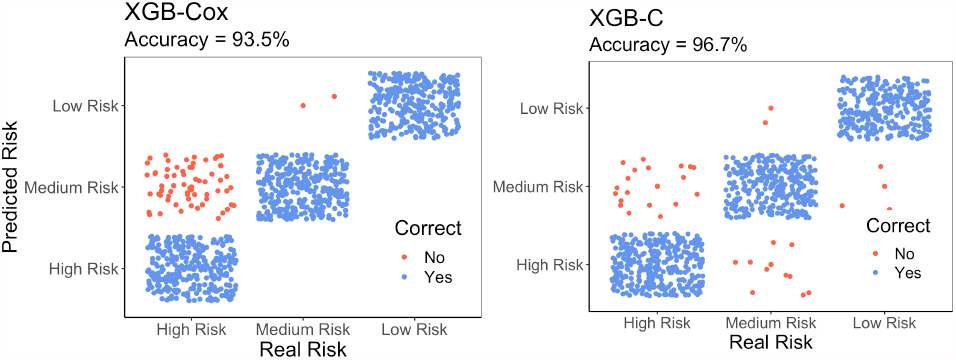
Survival calibration results from XGB-Cox (left panel) and XGB-C (right panel). The blue dots represent correctly classified (high/medium/low risk group) instances and red dots represent the misclassified instances.

### 3.4 Melanoma methylation dataset

In this section, we demonstrate the use of Xsurv through an analysis aiming to identify prognostic methylation biomarkers in melanoma. Matched methylation data (Illumina HumanMethylation 450K array) and gene expression from 470 melanoma samples were downloaded using the R package TCGA2STAT(Liu and Wan, 2015). All methylation values are arcsine transformed on beta values. For illustration purpose, we only select methylation CpG sites that have the strongest correlation (Spearman’s rho >0.3) with the gene expression level of the same gene. This shortlisted CpGs is closely related to the concept of cis-meQTL (Khan, 2018). This step resulted 9801 candidate CpGs as input data for prognostic biomarker discovery. To minimize patient heterogeneity, we focus the analysis on metastatic melanoma tumor sample. We further filtered 943 CpGs with small coefficient of variations (< 0.05). Stage is divided into two groups: stage I, II and I/II NOS into the low stage group and III, IV into the high stage group. The final data set contains 320 patients with 3 clinical covariates (including sex, age, stage) and 8858 CpGs. The prediction accuracy is evaluated by C-index and IBS. We compared the results of XGB and LGB with different models such as Cox model with Lasso method (Tibshirani, 1996) (Tibshirani, 1997), Random Forest (RF) (Ho, 1995) and GBM-Cox. The results are summarized in Table 2. We randomly resampled 100 times from the original dataset with a subsample of 80% of the samples (n=256 in each experiment). The average C-index and IBS in out-of-bag dataset were calculated. The top ranked CpG biomarkers are shown in Table 2. Among these CpGs, cg13629753 (in gene *GBP2*) and cg17209284 (in gene *SP140L*) were the most two selected by different models in Table 2 with 6 and 7 respectively. In addition, they are the only two that were selected by all Xsurv models. SP*140L* is a part of Speckled Protein (SP) family of chromatin ‘readers’ in humans and mice, and might play a key regulator role in silencing genes that establish immune cell identify and function (Fraschilla and Jeffrey, 2020). Another notable biomarker is cg11429292 (in gene *LAG3*). *LAG3* is a well-known immune checkpoint regulator mostly expressed in tumor infiltrated T cells and a promising target for immunotherapy in melanoma (Pardoll, 2012).We show the top 15 features selected by the LGB-Cox model who has the best C-index using the SHAP value plot (Lundberg and Lee, 2017) in Figure 3a and the aggregated feature selection results for LGB-Cox model in Figure 3b. Results for other models are represented in **Supplementary Fig. 3**. Since there are minor discrepancy for feature ranks between the SHAP- and gain-based results, we also provide the result based on gain in **Supplementary Fig. 2**. We present the recursive partitioning survival tree based on two most prognostic CpGs (cg13629753 and cg17209284) and stage in Figure 4. Through searching the Human Protein Atlas portal, we found that many top prognostic CpGs identified were in genes that are more pronounced in specific immune cells (https://www.proteinatlas.org/).Together, these results highlight the immunogenic characteristics of cutaneous melanoma and the underlying roles of these CpGs and genes warrant further investigation.

**Table 2.**
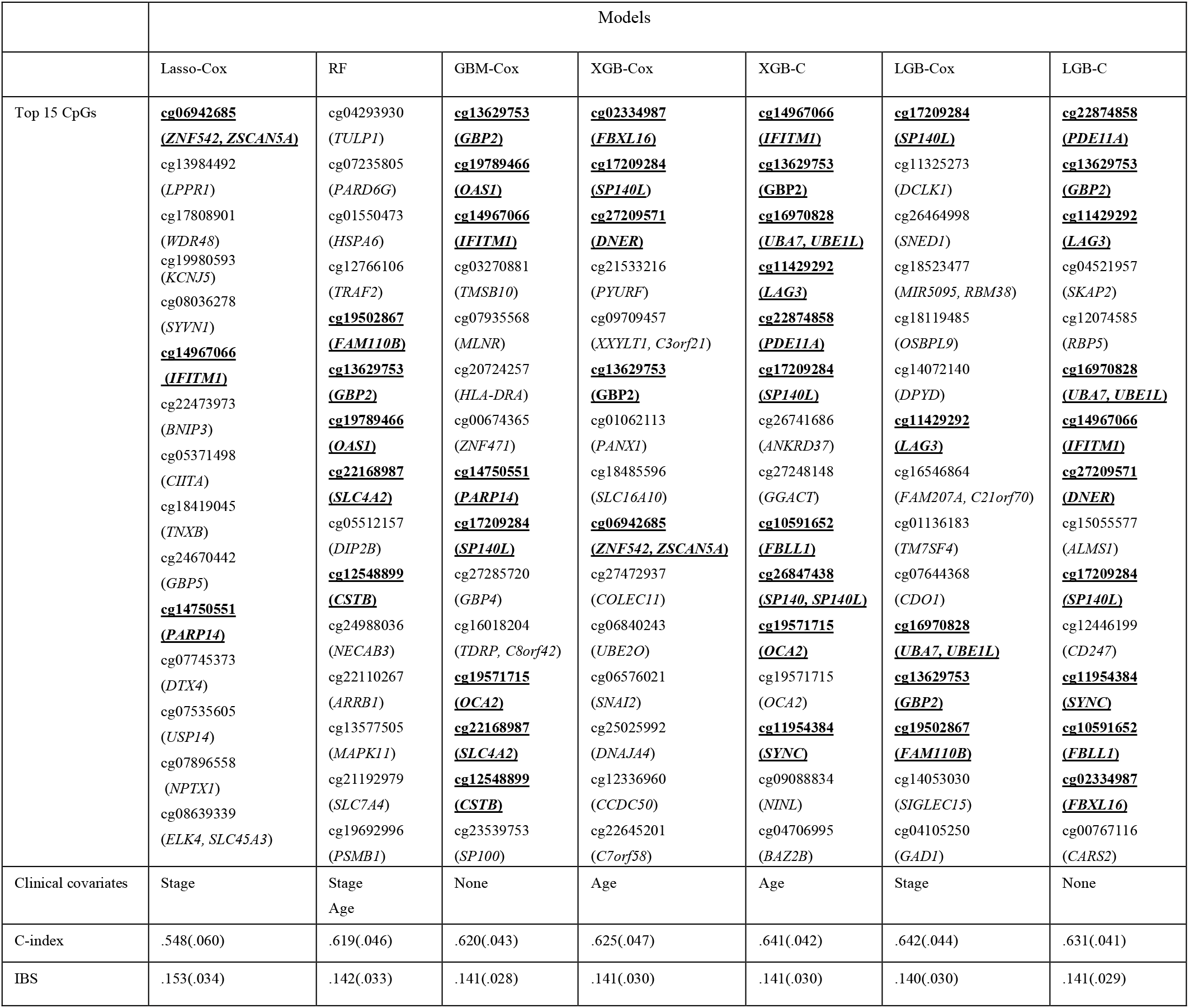
Top prognostic CpGs and corresponding genes selected by each method in the melanoma dataset. CpGs that were selected by multiple methods are highlighted.

**Fig.3.**
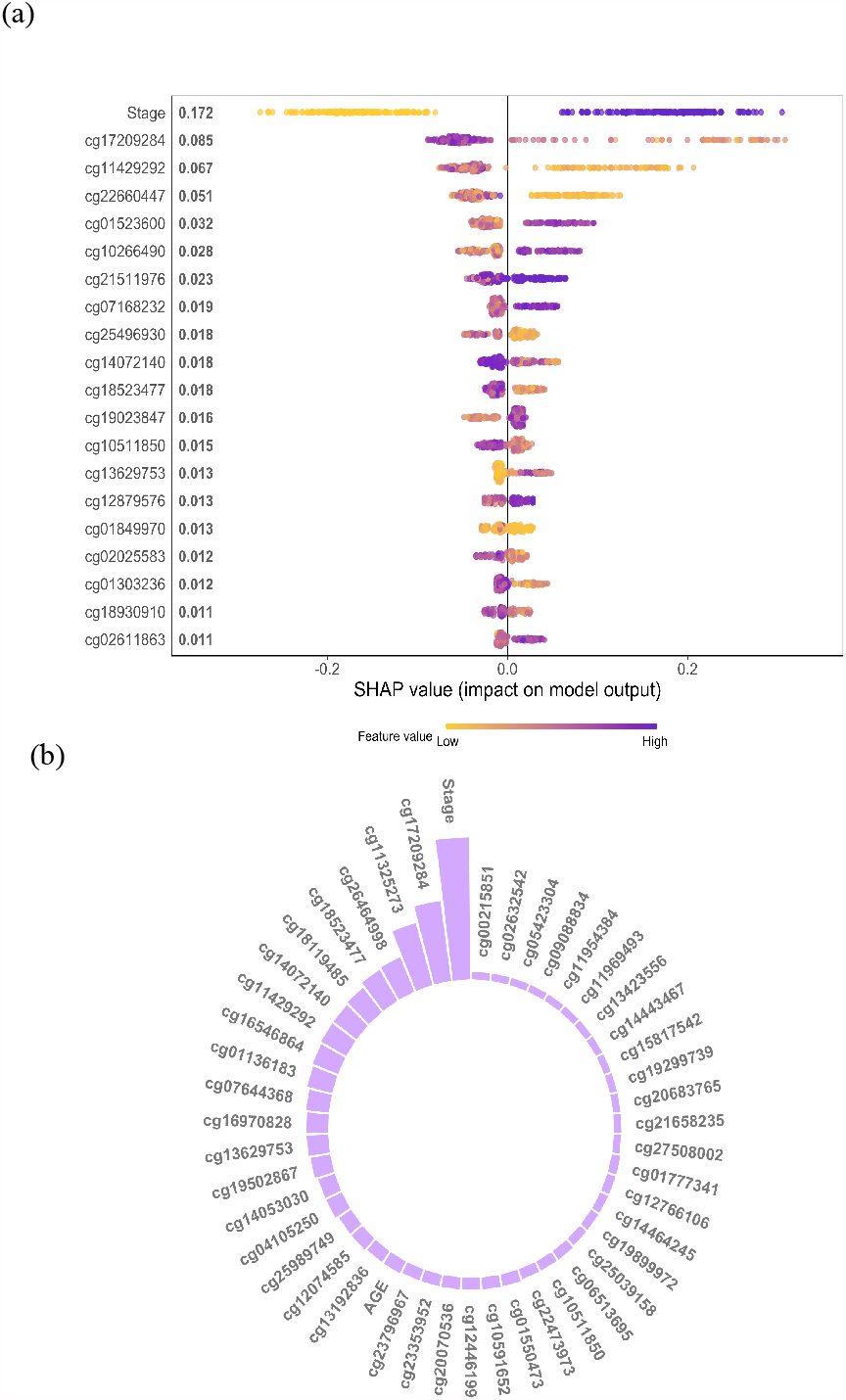
Top prognostic CpGs selected by XSurv (LGB-Cox) for predicting patient survival in melanoma. (a) SHAP summary plot of top 15 features from the LGB-Cox implementation. Each sample is represented by one dot at each row. (b) Circular barplot showing the frequency of the appearance of the CpG biomarkers in the top 15 features in each subsampling training (based on 100 replicates), where only those that appeared more than five times are displayed.

**Fig.4.**
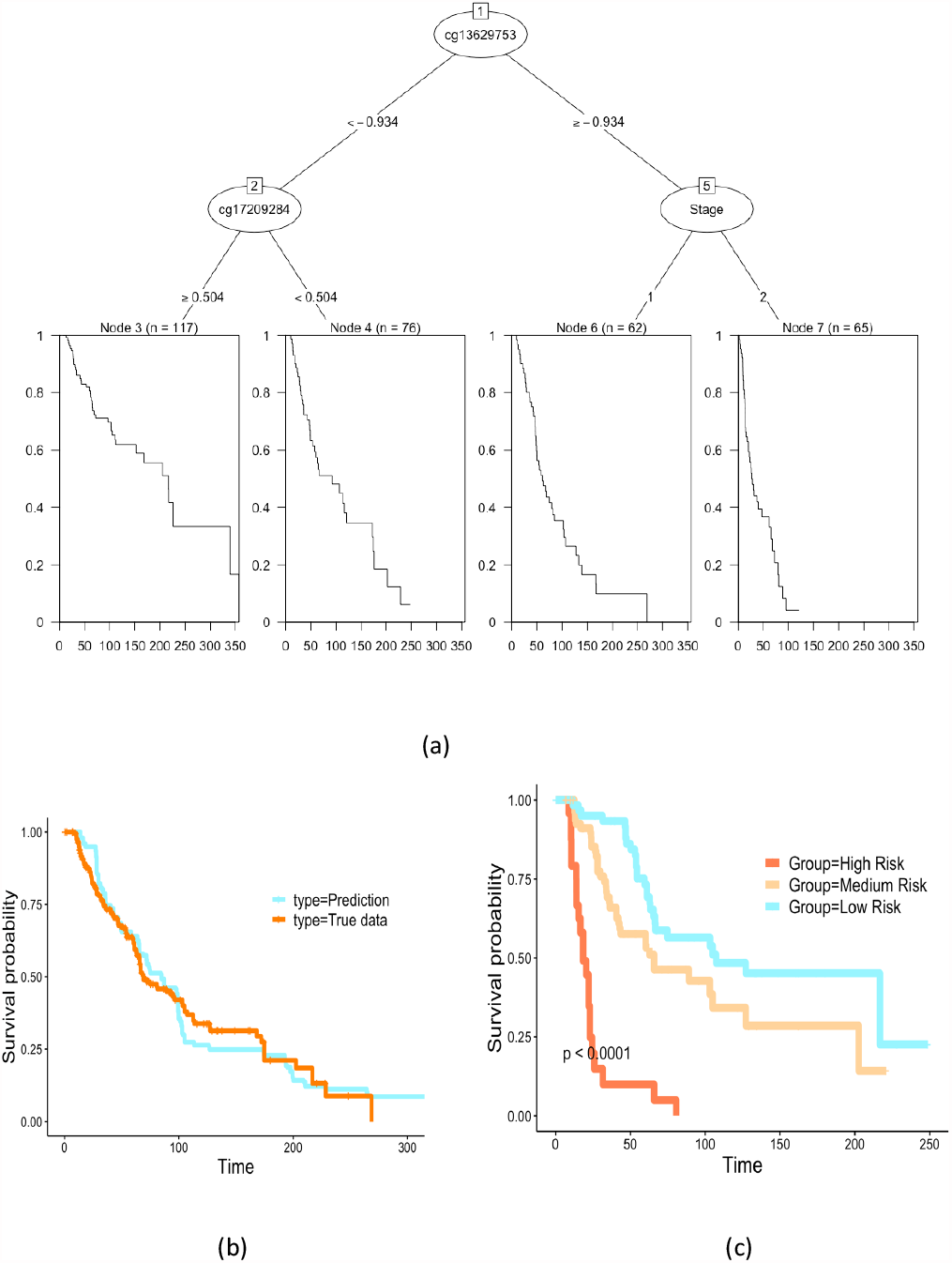
Melanoma patient survival prediction based on prognostic CpG biomarkers. (a) The recursive partitioning survival tree based on Stage and the top two robust prognostic CpGs in Table 2. (b) Kaplan-Meier plots comparing the predicted (based on LGB-Cox) and the observed patient overall survival data. (c) Kaplan-Meier plots comparing the patient subgroups stratified by the predicted risk group.

## 4 Conclusion

In this paper, we proposed a modern gradient boosting based framework, and a software package called **Xsurv**, to facilitate prognostic biomarker discovery with high-dimensional features and survival outcomes. **Xsurv** allows efficient survival analysis based on boosting either Cox PH loss or smoothed C-index, which relaxes the PH assumption in real data analyses. Our results suggested both XGB- and LGB-based approaches achieved considerably high prediction accuracy and robust biomarker selection under multiple scenarios considered in the simulation study. We recommend using LGB implementation when the data dimension is extremely high, due to its comparatively higher computationally efficiency over XGB. A distinctive feature of our package is that it allows survival time (subgroup) calibration and decision tree visualization, which will greatly aid in the interpretation of learned models. Finally, we applied the developed methods to a melanoma dataset to identify prognostic methylation biomarkers, and found many top predictive biomarkers to be indicative of potential immune regulations. Together, our **Xsurv** package provides a complementary and integrated tool for biomarker discovery with high-dimensional molecular, clinical or imaging data. Future efforts will be needed to allow the package to handle competing risks and to explore putative mediation effects.

## Acknowledgements

The authors would like to thank Dr. Hongzhe Li for helpful discussion many years ago that shaped the early draft of this paper.

## References

1. Brier, G.W. VERIFICATION OF FORECASTS EXPRESSED IN TERMS OF PROBABILITY. Monthly Weather Review 1950;78(1):1–3.

2. Chen, T. and Guestrin, C. XGBoost: A Scalable Tree Boosting System. In, Proceedings of the 22nd ACM SIGKDD International Conference on Knowledge Discovery and Data Mining. San Francisco, California, USA: Association for Computing Machinery; 2016. p. 785–794.

3. Chen, T., et al. 2021. Xgboost: Extreme Gradient Boosting. https://github.com/dmlc/xgboost

4. Chen, Y., et al. A Gradient Boosting Algorithm for Survival Analysis via Direct Optimization of Concordance Index. Computational and Mathematical Methods in Medicine 2013;2013:873595.

5. Cox, D.R. Regression Models and Life-Tables. Journal of the Royal Statistical Society. Series B (Methodological) 1972;34(2):187–220.

6. Draper, N. and Smith, H. Applied Regression Analysis, 2d Edition. New York: John Wiley & Sons, Inc.; 1981.

7. Efroymson, M.A. Multiple regression analysis. Mathematical Methods for Digital Computers, Wiley, New York. 1960.

8. Fraschilla, I. and Jeffrey, K.L. The Speckled Protein (SP) Family: Immunity’s Chromatin Readers. Trends Immunol 2020;41(7):572–585.

9. Freund, Y. and Schapire, R.E. A Decision-Theoretic Generalization of On-Line Learning and an Application to Boosting. Journal of Computer and System Sciences 1997;55(1):119–139.

10. Friedman, J.H. Greedy function approximation: A gradient boosting machine. Ann. Statist. 2001;29(5):1189–1232.

11. Greenwell, B., et al. 2007. Generailzed Boosted Models: A guide to the gbm package. https://CRAN.R-project.org/package=gbm

12. Harrell, F.E., et al. Evaluating the yield of medical tests. JAMA 1982;247(18):2543–2546.

13. Hastie, T., Tibshirani, R. and Friedman, J.H. Boosting and Additive Trees. 2009. Ho, T.K. Random decision forests. Proceedings of 3rd International Conference on Document Analysis and Recognition 1995;1:278–282.

14. Hocking, R.R. The Analysis and Selection of Variables in Linear Regression. Biometrics 1976.

15. Jeschke, J., et al. DNA methylation–based immune response signature improves patient diagnosis in multiple cancers. The Journal of clinical investigation 2017;127(8):3090–3102.

16. Khan, S. Advances in Usage of Venom Proteins as Diagnostics and Therapeutic Mediators. Protein Pept Lett 2018;25(7):610–611.

17. Kuhn, M. 2020. caret:Classification and Regression Training. https://CRAN.R-project.org/package=caret

18. Li, H. and Luan, Y. Boosting proportional hazards models using smoothing splines, with applications to high-dimensional microarray data. Bioinformatics 2005;21(10):2403–2409.

19. Liu, Z. and Wan, Y.-W. 2015. TCGA2STAT: Simple TCGA Data Access for Integrated Statistical Analysis in R. http://www.liuzlab.org/TCGA2STAT/

20. Lundberg, S. and Lee, S.-l. A unified approach to interpreting model predictions. In.; 2017.

21. Mason, L., et al. Boosting Algorithms as Gradient Descent in Function Space. Advances in Neural Information Processing Systems 12.MIT Press 1999:512–518.

22. Pardoll, D.M. The blockade of immune checkpoints in cancer immunotherapy. Nat Rev Cancer 2012;12(4):252–264.

23. Simon, N., et al. A Sparse-Group Lasso. Journal of Computational and Graphical Statistics 2013;22(2):231–245.

24. Tibshirani, R. Regression Shrinkage and Selection Via the Lasso. Journal of the Royal Statistical Society: Series B (Methodological) 1996;58(1):267–288.

25. Tibshirani, R. The Lasso method for variable selection in the Cox model. Statistics in Medicine 1997;16(4):385–395.

26. Yuan, M. and Lin, Y. Model selection and estimation in regression with grouped variables. Journal of the Royal Statistical Society: Series B (Statistical Methodology) 2006;68(1):49–67.

