## Supplemental Materials for "Efficient gradient boosting for prognostic biomarker discovery"

---

### 1. Second derivative for Cox model

Proportional hazard model was introduced by Cox, with the assumption that the features or covariates are multiplicatively related to the hazard, that is

$$\lambda(t|x,\theta) = \lambda_0(t)\exp(\boldsymbol{x}^{\boldsymbol{T}}\theta)$$

Under this proportional hazard assumption, we don't need to optimize the likelihood function, instead, we just need to optimize the Cox partial likelihood function

$$L_p(\theta) = \prod_{i\in E}\frac{\exp\ (\boldsymbol{x}_i^{\boldsymbol{T}}\theta)}{\sum_{j:t_j\geq t_i}\exp\ (\boldsymbol{x}_j^{\boldsymbol{T}}\theta)}$$

where $E$ is the set of data that events happen.

We use the partial likelihood as the loss function in XGB or LGB:

$$L(y,F) = -\sum_{i=1}^{n}\delta_i\left\{H(x_i) - \log\left(\sum_{j:t_j\geq t_i}\mathrm{H}(\mathrm{x_j})\right)\right\}$$

The first derivative is

$$\frac{\partial L(y,F)}{\partial H(x_i)} = -\frac{\partial}{\partial H(x_i)}\delta_i\left\{H(x_i) - \log\left(\sum_{j:t_j\geq t_i}\exp\left(H(x_j)\right)\right)\right\}$$

where $\tilde{\pi}_{ij}$ is defined in 2.1.1.

The second derivative is

$$\frac{\partial^2 L(y,F)}{\partial H(x_i)^2} = -\frac{\partial}{\partial H(x_i)} \sum_k \delta_k \frac{I(t_i \geq t_k) \exp\left(H(x_j)\right)}{\sum_j I\left(t_j \geq t_k\right) \exp\left(H(x_j)\right)}$$

$$= \sum_k \delta_k \frac{I(t_i \geq t_k) \exp\left(H(x_j)\right)}{\sum_j I\left(t_j \geq t_k\right) \exp\left(H(x_j)\right)} - \sum_k \delta_k \left(\frac{I(t_i \geq t_k) \exp\left(H(x_j)\right)}{\sum_j I\left(t_j \geq t_k\right) \exp\left(H(x_j)\right)}\right)^2$$

$$= -\sum_j \delta_j \tilde{\pi}_{ij} \left(1 - \tilde{\pi}_{ij}\right).$$

### 2. SCI implement

The second derivative of C-index loss function is given by

$$h_i = \frac{\partial g_i}{\partial H(x_i)} = \frac{\alpha^2}{|\mathcal{P}|} \left\{ \sum_{(i,j)\in\mathcal{P}} \frac{-\exp\left(\alpha\left(H(x_i) - H\left(x_j\right)\right)\right) * (1 - \exp\left(\alpha\left(H(x_i) - H\left(x_j\right)\right)\right))}{\left[1 + \exp\left(\alpha\left(H(x_i) - H\left(x_j\right)\right)\right)\right]^3} \right\} +$$

$$\frac{\alpha^2}{|\mathcal{P}|} \left\{ \sum_{(k,i)\in\mathcal{P}} \frac{-\exp\left(\alpha\left(H(x_k) - H(x_i)\right)\right) * (1 - \exp\left(\alpha\left(H(x_k) - H(x_i)\right)\right))}{\left[1 + \exp\left(\alpha\left(H(x_k) - H(x_i)\right)\right)\right]^3} \right\}$$
