## Supplemental Figures for "Efficient gradient boosting for prognostic biomarker discovery"

---

### 1. Additional Simulation Information

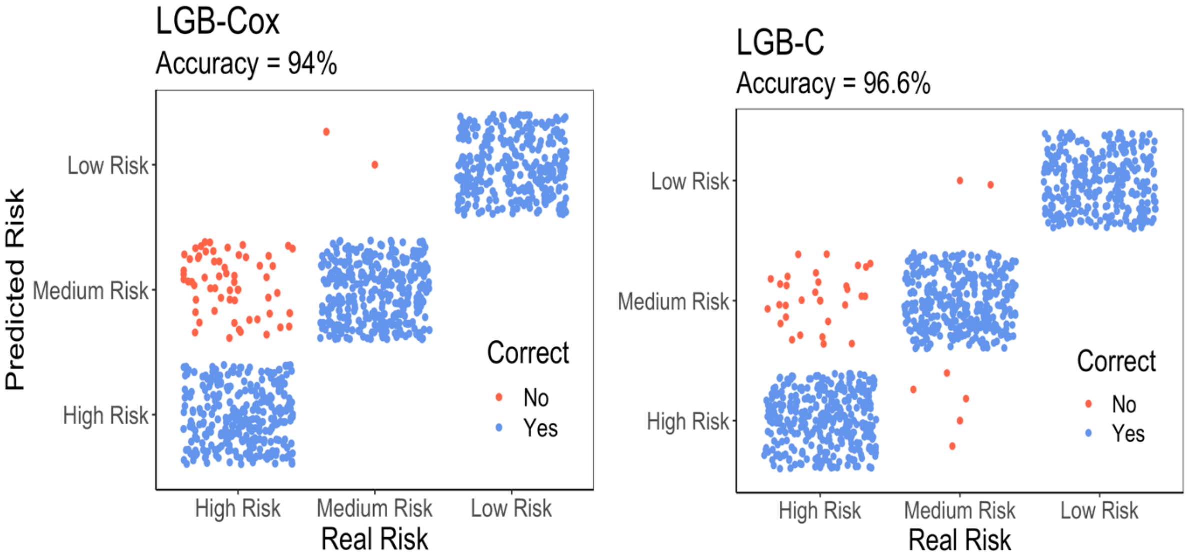


**Supplementary Fig.1**
Survival outcome calibration results from LGB-Cox (left panel) and LGB-C (right panel). The blue dots represent correctly classified (high/medium/low risk group) instances and red dots represent the misclassified instances.

### 2. Extra Real data implement

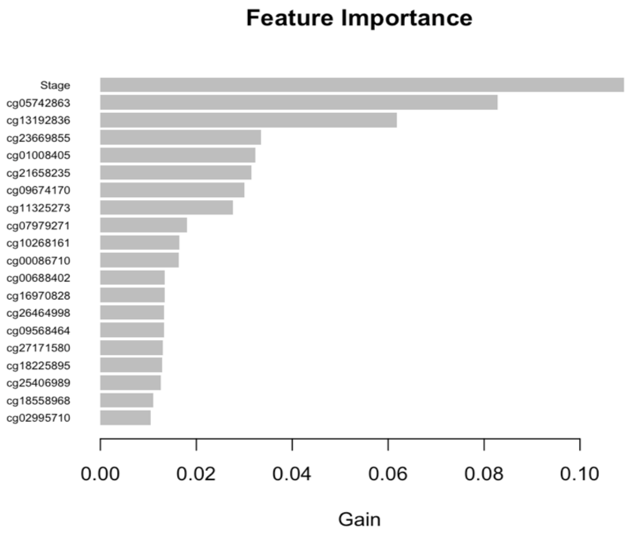


**Supplementary Fig.2**
Top 15 prognostic features for selected LGB-Cox model by gain.

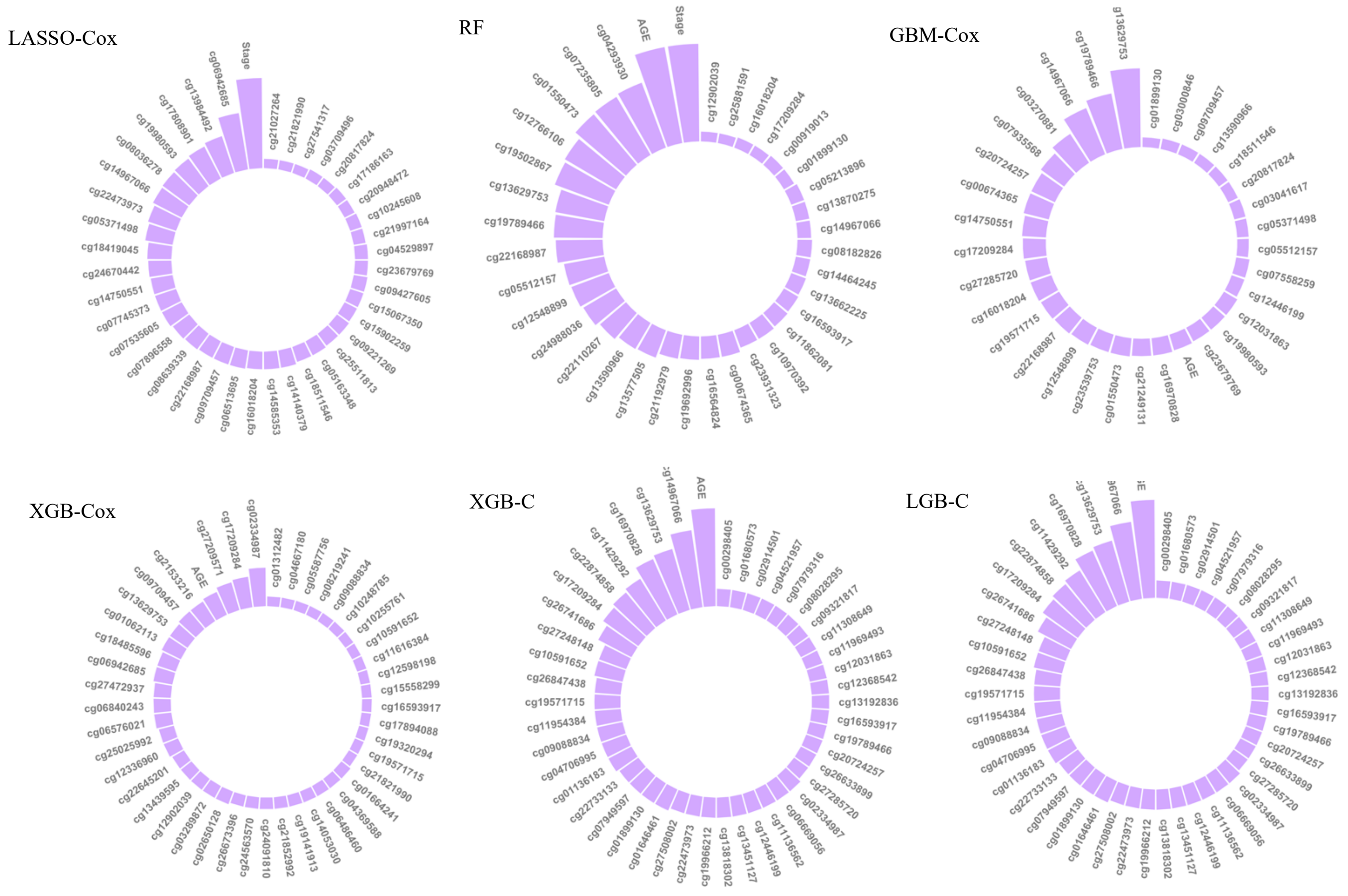


**Supplementary Fig.3**
Circular barplot showing the frequency of the appearance of the CpG biomarkers in the top 15 features in each subsampling training (based on 100 replicates) where only those that appeared more than five times are displayed from different models as labeled.

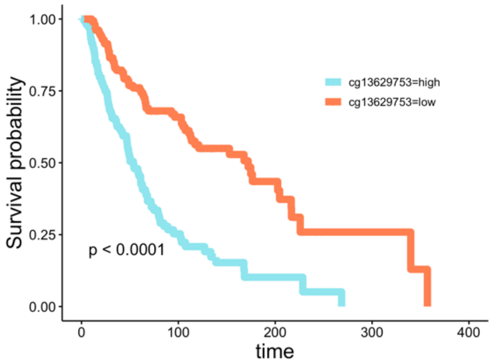


(a)

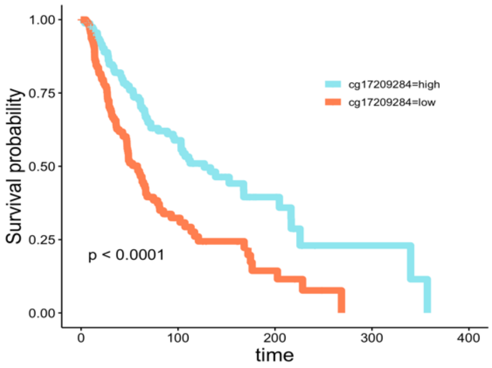


(b)

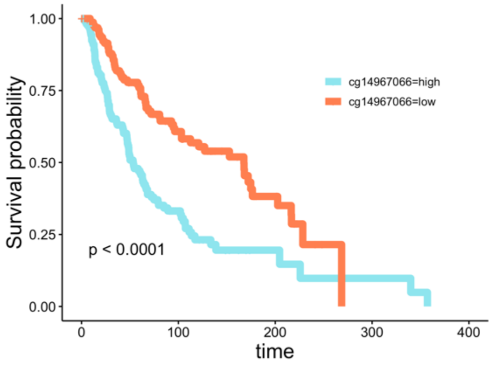


(c)

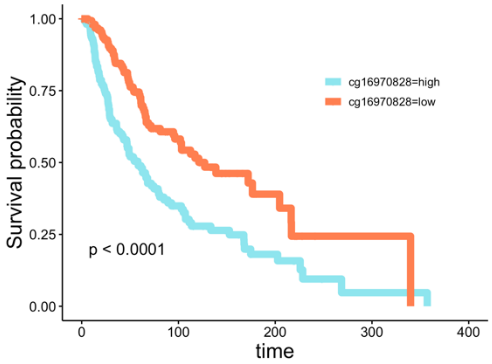


(d)

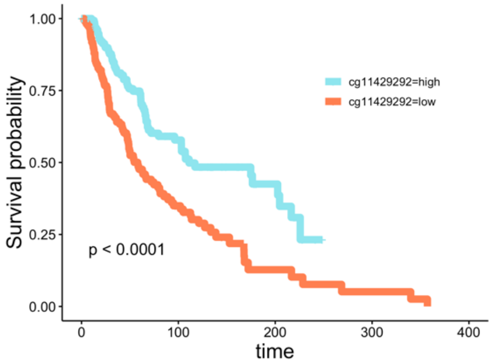


(e)

**Supplementary Fig.4**

Kaplan-Meier curves comparing the patient subgroups stratified by the underlined CpGs that were selected by different models at least three times in Table 2.

(a) cg13629753, (b) cg17209284, (c) cg14967066, (d) cg16970828, (e) cg11429292.
